## Supplemental Material for "Aminergic and peptidergic modulation of Insulin-Producing Cells in *Drosophila*"

### Supplements

**Table S1: Abbreviations used for receptors, neuropeptides, and biogenic amines.**

| Abbreviation | Full name |
|---|---|
| **Biogenic amine receptors** | |
| 5-HT1A | 5-Hydroxytryptamine receptor 1A |
| 5-HT1B | 5-Hydroxytryptamine receptor 1B |
| 5-HT2A | 5-Hydroxytryptamine receptor 2A |
| 5-HT2B | 5-Hydroxytryptamine receptor 2B |
| 5-HT7 | 5-Hydroxytryptamine receptor 7 |
| CG13579 | Orphan |
| Dop1R1 | Dopamine 1-like receptor 1 |
| Dop1R2 | Dopamine 1-like receptor 2 |
| Dop2R | Dopamine 2-like receptor |
| DopEcR | Dopamine Ecdysone receptor |
| Oamb | Octopamine receptor in mushroom bodies |
| Octalpha2R | α2-adrenergic-like octopamine receptor |
| Octbeta1R | Octopamine beta 1 receptor |
| Octbeta2R | Octopamine beta 2 receptor |
| Octbeta3R | Octopamine beta 3 receptor |
| Oct-TyR | Octopamine-Tyramine receptor |
| TyR | Tyramine receptor |
| **Neuropeptide receptors** | |
| AstA-R1 | Allatostatin A receptor 1 |
| AstA-R2 | Allatostatin A receptor 2 |
| CCAP-R | Crustacean cardioactive peptide receptor |
| CCHa2-R | CCHamide-2 receptor |
| CG10738 | Orphan |
| CNMaR | CNMamide receptor |
| Dh31-R | Diuretic hormone 31 receptor |
| Dh44-R1 | Diuretic hormone 44 receptor 1 |
| Dh44-R2 | Diuretic hormone 44 receptor 2 |
| FMRFaR | FMRFamide receptor |
| hec | hector |
| InR | Insulin-like receptor |
| Lkr | Leucokinin receptor |
| NPFR | Neuropeptide F receptor |
| Pdfr | Pigment-dispersing factor receptor |
| rk | rickets |
| RYa-R | RYamide receptor |
| SIFaR | SIFamide receptor |
| sNPF-R | short Neuropeptide F receptor |
| SPR | Sex peptide receptor |
| TkR86C | Tackykinin-like receptor at 86C |
| TrissinR | Trissin receptor |
| **Biogenic amines** | |
| 5-HT | 5-Hydroxytryptamine (serotonin) |
| DA | Dopamine |
| OA | Octopamine |
| Tyr | Tyramine |
| **Neuropeptides** | |
| AstA | Allatostatin-A |
| DH31 | Diuretic hormone 31 |
| LK | Leucokinin |
| MS | Myosuppressin |
| sNPF | short Neuropeptide F |
| TK | Tachykinin |

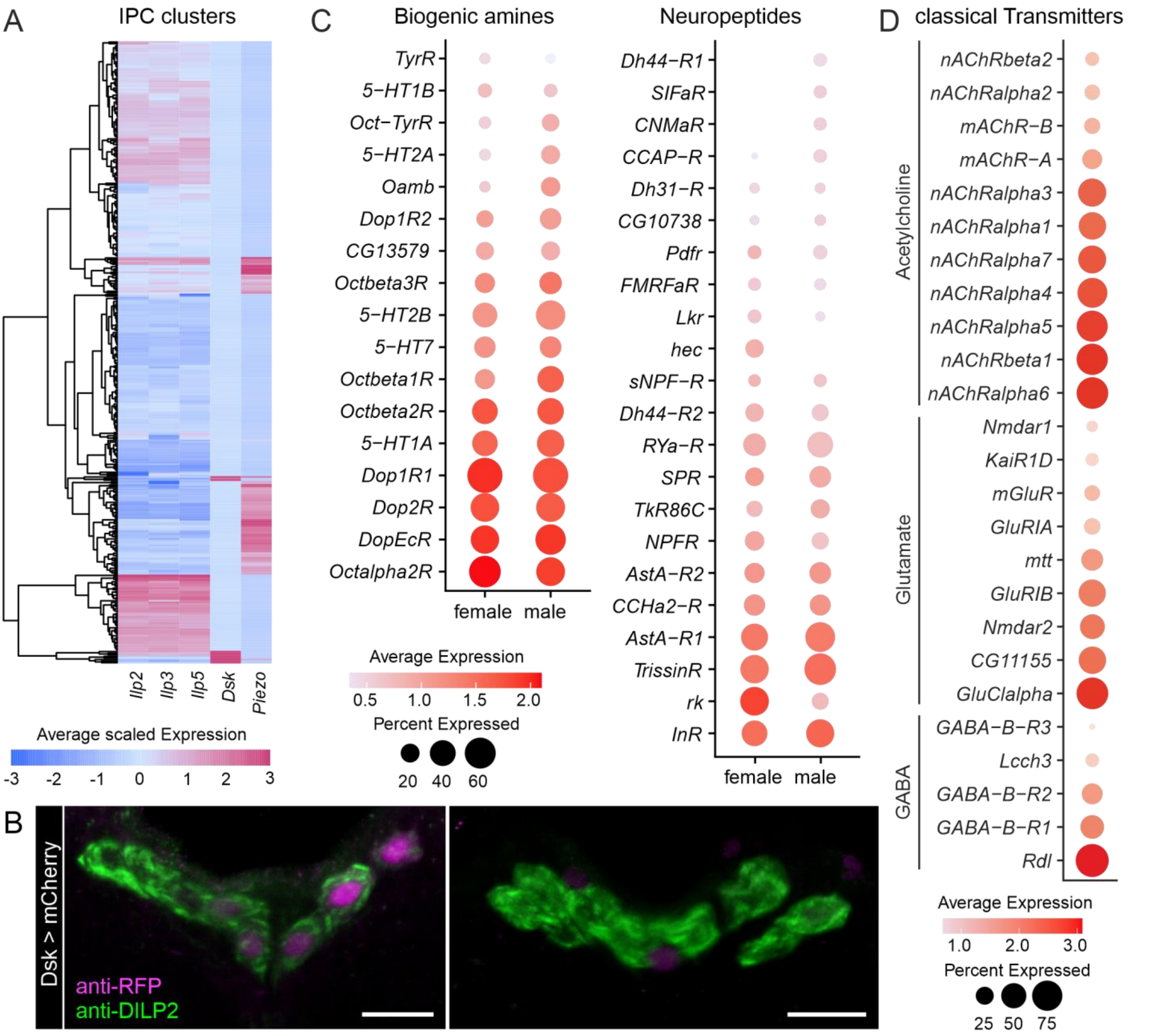


**Figure S1: IPCs express receptors for biogenic amines, neuropeptides, and classical neurotransmitters.. A:** Cluster analysis of IPCs based on *Ilp2*, *Ilp3*, *Ilp5*, *Dsk*, and *Piezo* expression. Expression levels are depicted as a heatmap where each row represents a single cell. Expression was scaled based on all genes in the dataset. Therefore, negative values indicate low expression. **B:** Representative confocal stacks from two brains showing sparse *Dsk-T2A-GAL4* driven mCherry expression (amplified with RFP antibody and shown in magenta) in IPCs (labelled with DILP2 antibody and shown in green). Scale bars = 10 µm. **C:** Comparison of biogenic amine and neuropeptide receptor expression in IPC transcriptomes derived from males and females. **D:** Expression of classical fast-acting neurotransmitter receptors in IPC transcriptomes.

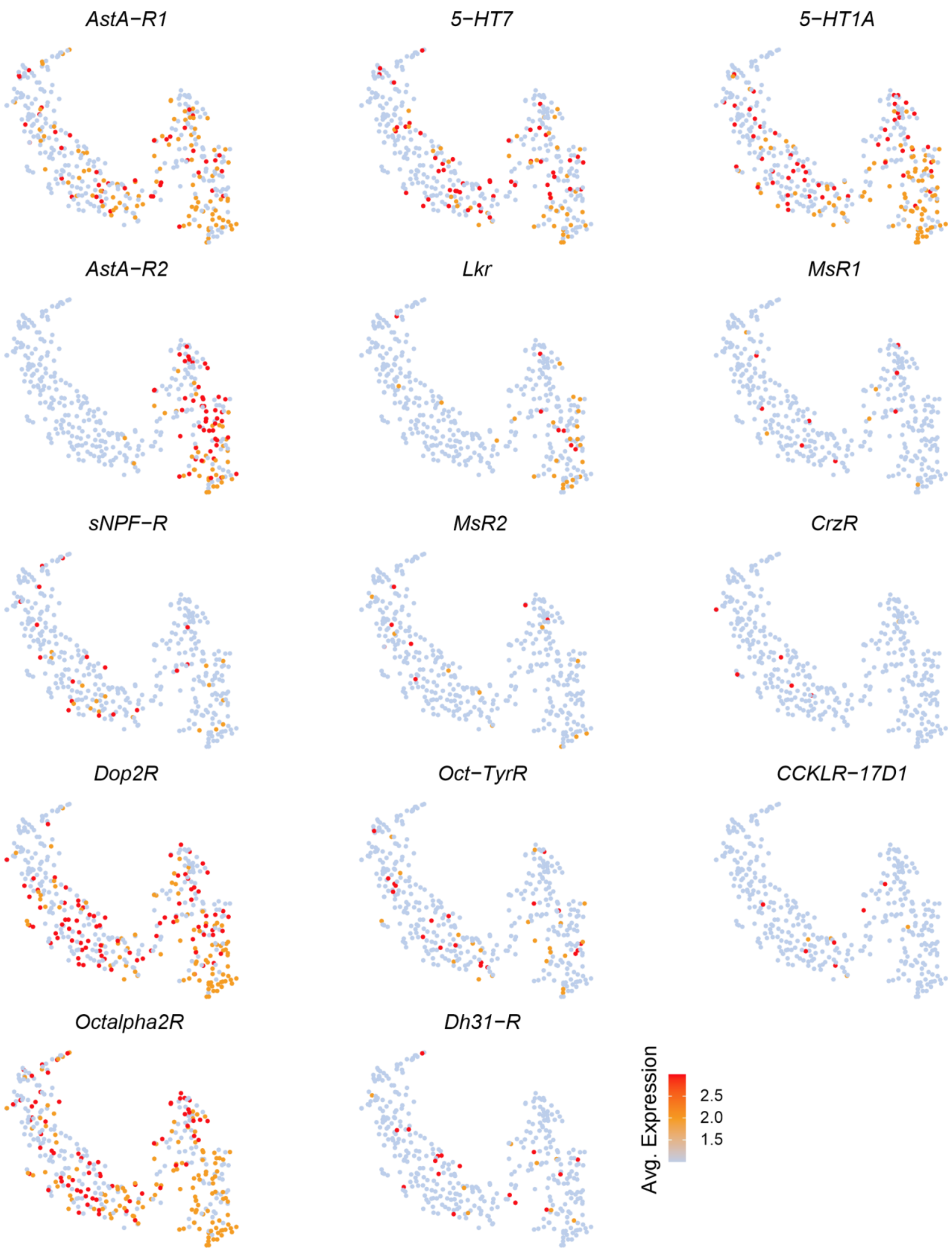


**Figure S2: Expression of select neuromodulator receptors across individual IPCs**. t-SNE plots showing expression of select biogenic amine and neuropeptide receptors across IPC transcriptomes. Note that some receptors such as *AstA-R1* and *5-HT1A* are broadly expressed whereas others, for example *MsR1* and *MsR2*, are sparsely expressed.

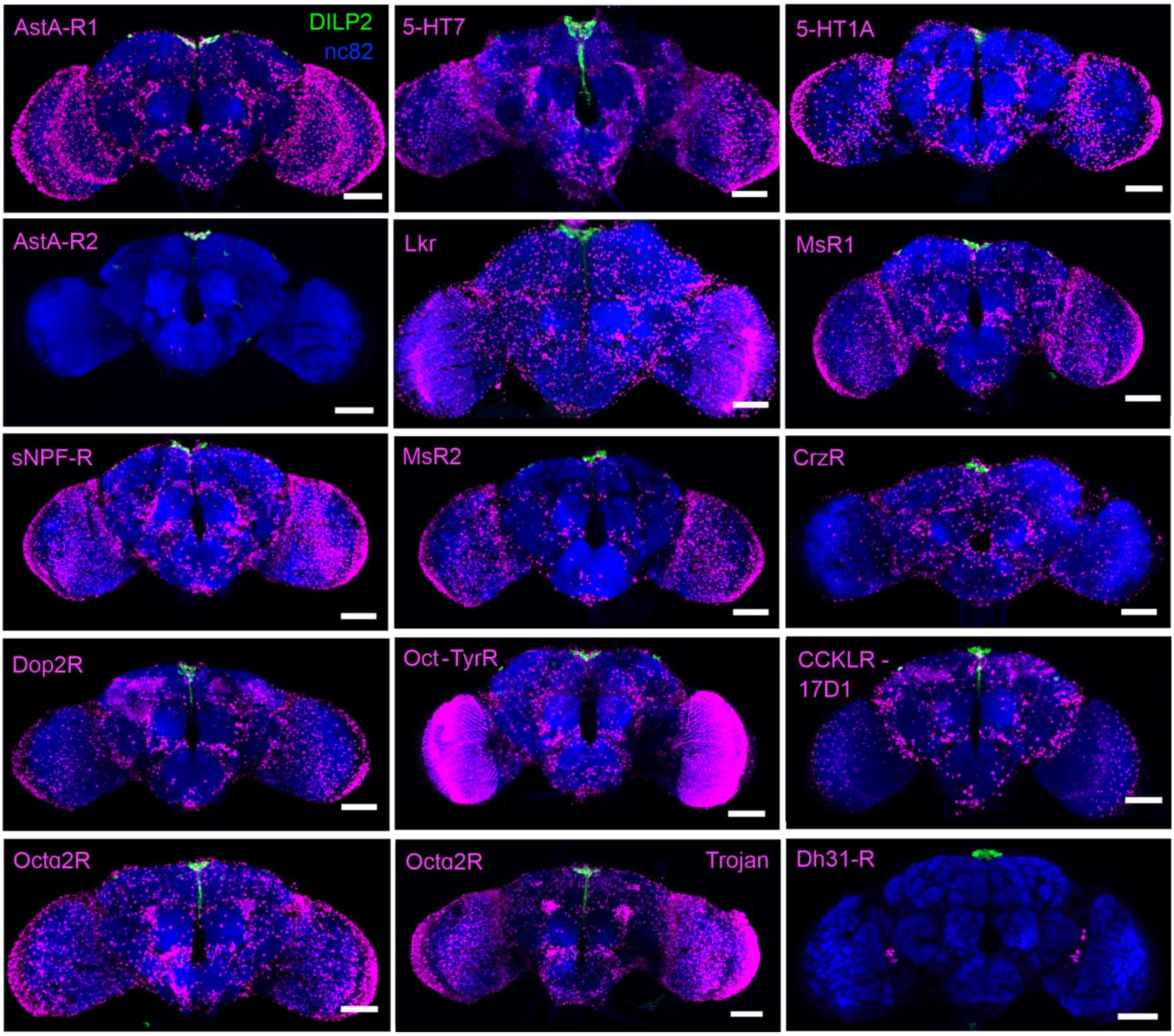


**Figure S3: Overview of neuromodulator receptor expression in the brain.** T2A-GAL4 knock-in lines for different receptors drive broad expression of nuclear mCherry (magenta). IPCs (green) and the neuropil (blue) have been labelled using DILP2 and nc82 antibodies, respectively. Note that AstAR2 is mainly expressed in the IPCs. For Octα2R, an independent GAL4 (Trojan) was also tested. Scale bars = 50 µm.

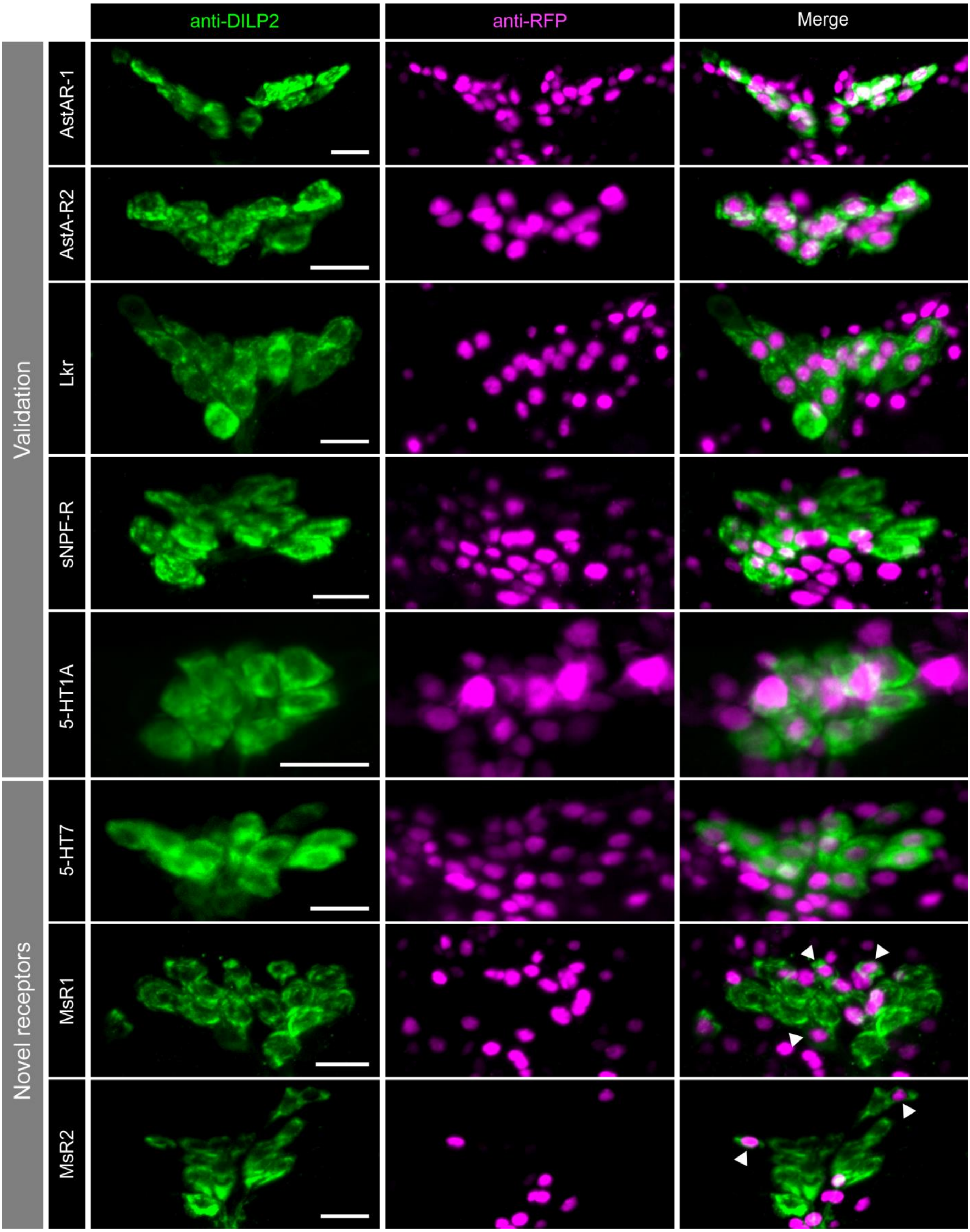


**Figure S4: Neuromodulator receptors expressed in IPCs.** Representative confocal stacks showing expression of different neuromodulator receptors (magenta) in IPCs (labelled using DILP2 antibody in green). Receptor expression was assessed by driving nuclear mCherry reporter using -T2A-GAL4 knockin lines for allatostatin-A receptor 1 (AstA-R1), allatostatin-A receptor 2 (AstA-R2), leucokinin receptor (Lkr), short neuropeptide F receptor (sNPF-R), 5-hydroxytryptamine (5-HT) receptor 1A (5-HT1A), 5-HT receptor 7 (5-HT7), myosuppressin receptor 1 (MsR1) and myosuppressin receptor 2 (MsR2). Scale bars = 10 µm.

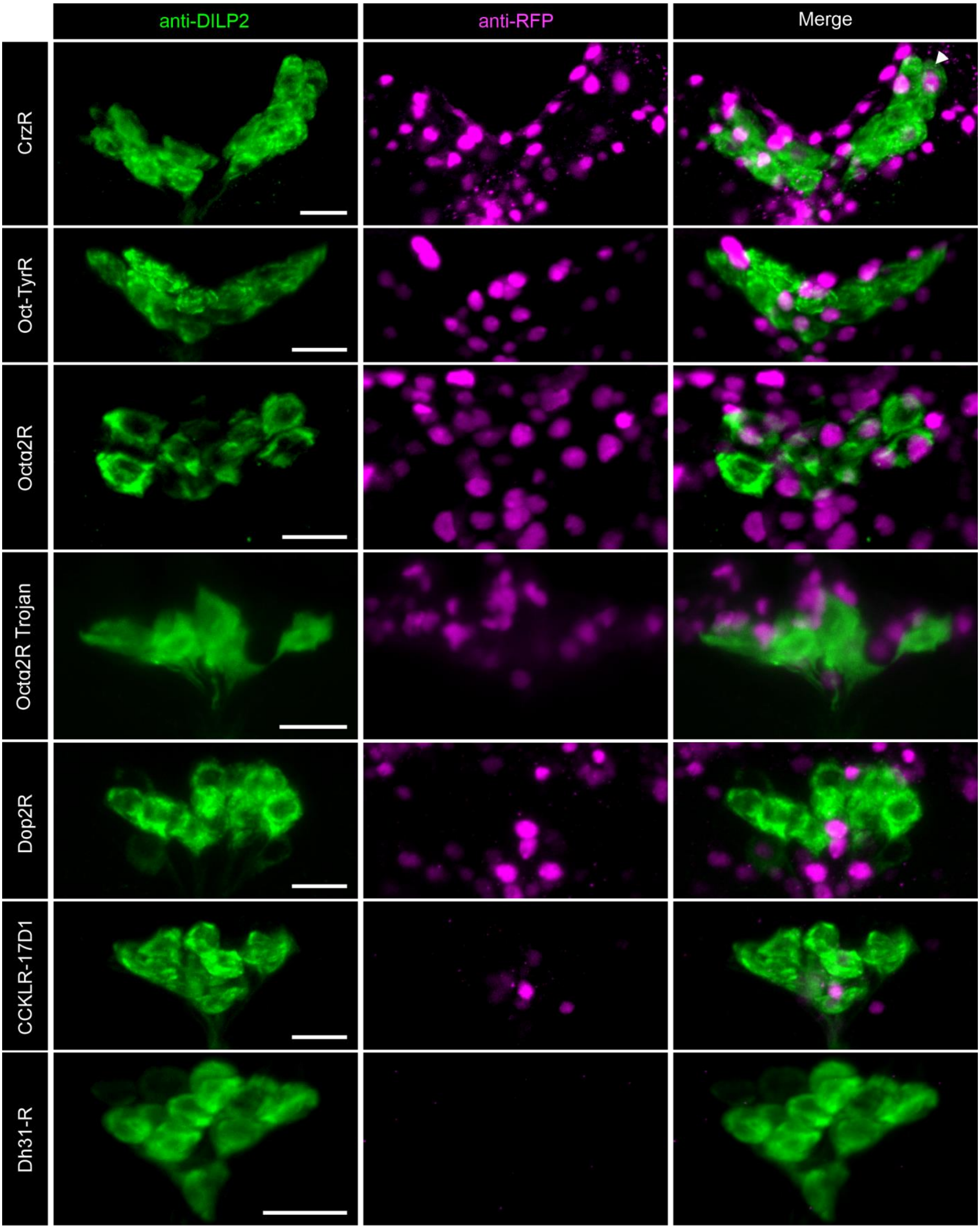


**Figure S5: Neuromodulator receptors not expressed in IPCs.** Representative confocal stacks showing a lack of expression of different neuromodulator receptors (magenta) in IPCs (labelled using DILP2 antibody in green). Receptor expression was assessed by driving nuclear mCherry reporter using GAL4 lines for corazonin receptor (CrzR), octopamine (Oct)-tyramine receptor (Oct-TyrR), Oct alpha receptor 2 (Octα2R), Octα2R Trojan, dopamine receptor 2 (Dop2R), cholecystokinin-like receptor at 17D1 (CCKLR-17D1) and diuretic hormone 31 receptor (Dh31-R). Scale bars = 10 µm. With the exception of Octα2R Trojan, all other lines were T2A-GAL4 knock-ins.

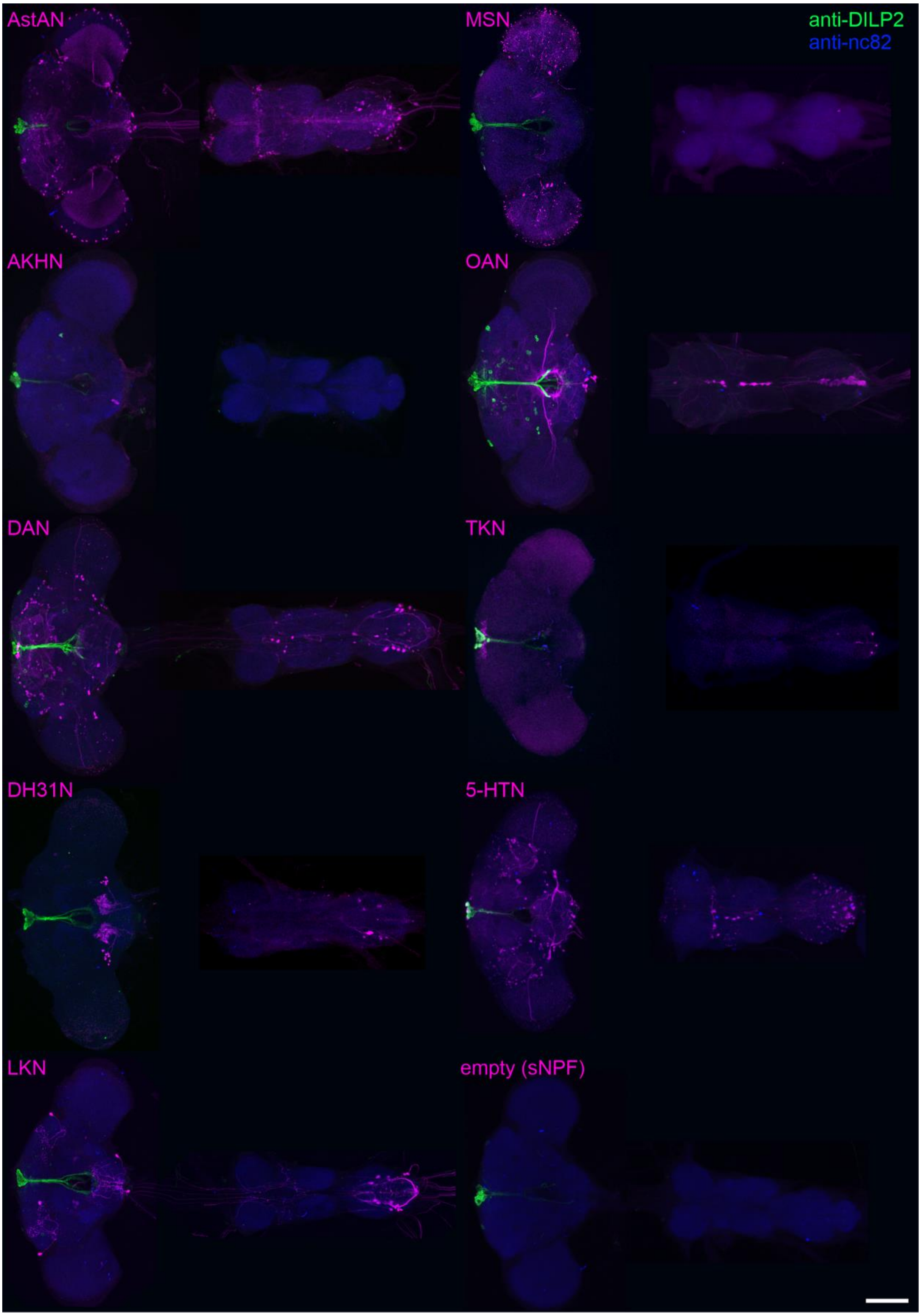


**Figure S6: Expression of the ModN driver lines in the brain and VNC. The** expression pattern of each ModN line was visualized via GFP-expression (magenta) in the fly brain and VNC using the respective GAL4 driver line. IPCs were visualized using anti-DILP2 (green), and anti-nc82 immunohistochemical labeling was used to stain the neuropil (blue). The sNPFN driver line was expected to drive expression in sNPF-expressing neurons. However, anatomical labeling revealed no expression in the central nervous system across all inspected animals. Therefore, the sNPF driver lines was “empty” and served as a negative control. Scale bar = 100 µm.

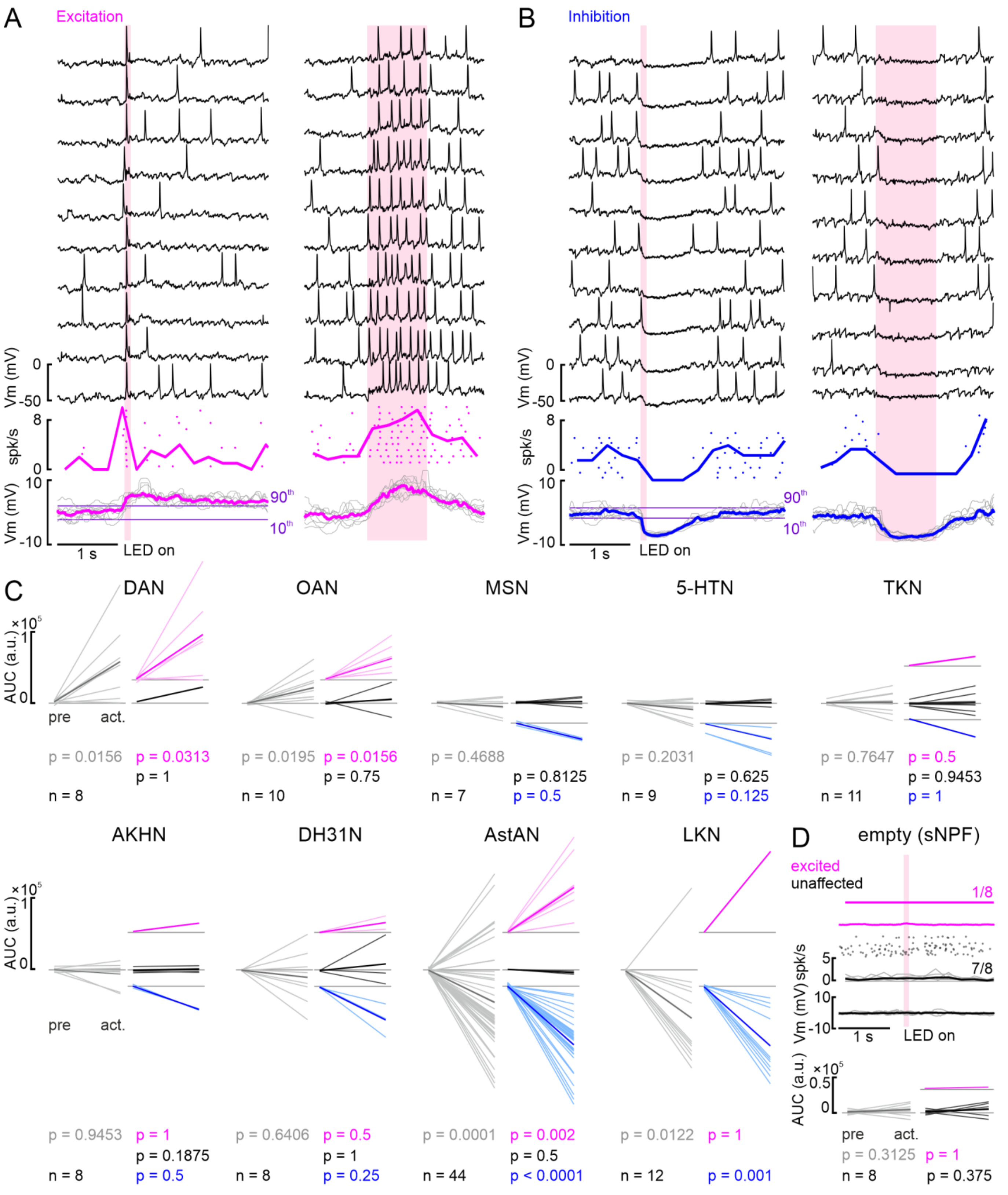


**Figure S7: IPC responses to repeated activation of ModNs and statistical analysis of IPC responses to ModN activation for all lines. A:** Example recording from IPC shows excitation upon DAN activation across ten repetitions (black traces) for 100 ms activation (left) and 1 s activation (right). Spike events are displayed as dots, with each row representing one repetition of the activation, and the spike frequency as thick, color-coded line. The membrane potential of each repetition is displayed in gray with the median as thick, color-coded line. Purple lines indicate the upper and lower thresholds for the cluster analysis. **B:** Recording of example IPC that shows inhibition upon AstAN activation. Plot details as in A. **C:** AUC values of individual IPCs averaged in a 1 s window before activation onset (pre) and in a window from activation onset to 1 s after activation offset (act.) for all ModN lines. Grey, all recorded IPCs, colors, three clusters (color-coded as before). P-values were calculated using a paired Wilcoxon signed-rank test. **D:** Upper panel: IPC responses to activation of the ‘empty’ sNPF line. One IPC showed a slight membrane potential increase that did not cause spikes but crossed the upper threshold, and 7 IPCs remained completely unaffected. Lower panel: averaged AUC values of the 8 IPCs recorded during activation of the ‘empty’ line (details as in C). Activation of this ‘empty’ line had no significant effect on the IPCs, as expected.

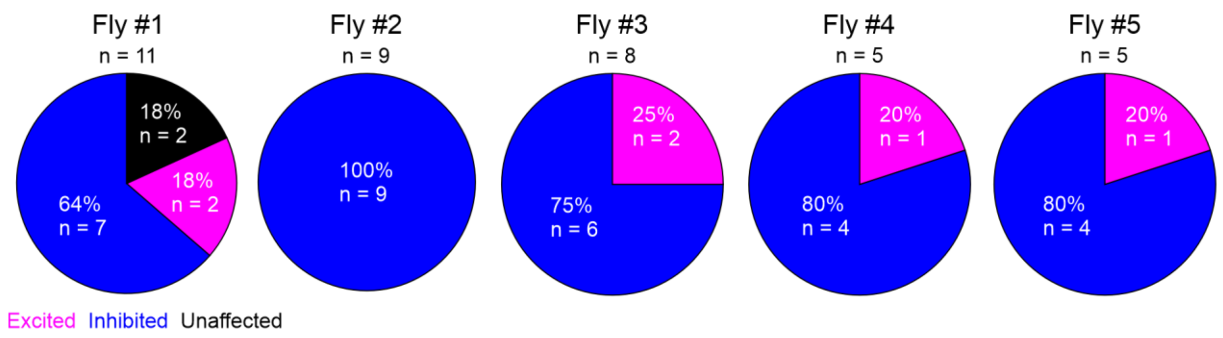


**Figure S8: Cluster distribution of multiple IPCs recorded in the same individuals during AstANs activation.** The recorded IPCs in Fly #1 (example from Figure 3) were divided into an excited, inhibited, and an unaffected cluster based on threshold clustering. In Fly #2, all IPCs were inhibited, while in Fly #3 – Fly #5 the recorded IPCs were either inhibited or excited. n = number of recorded IPCs.

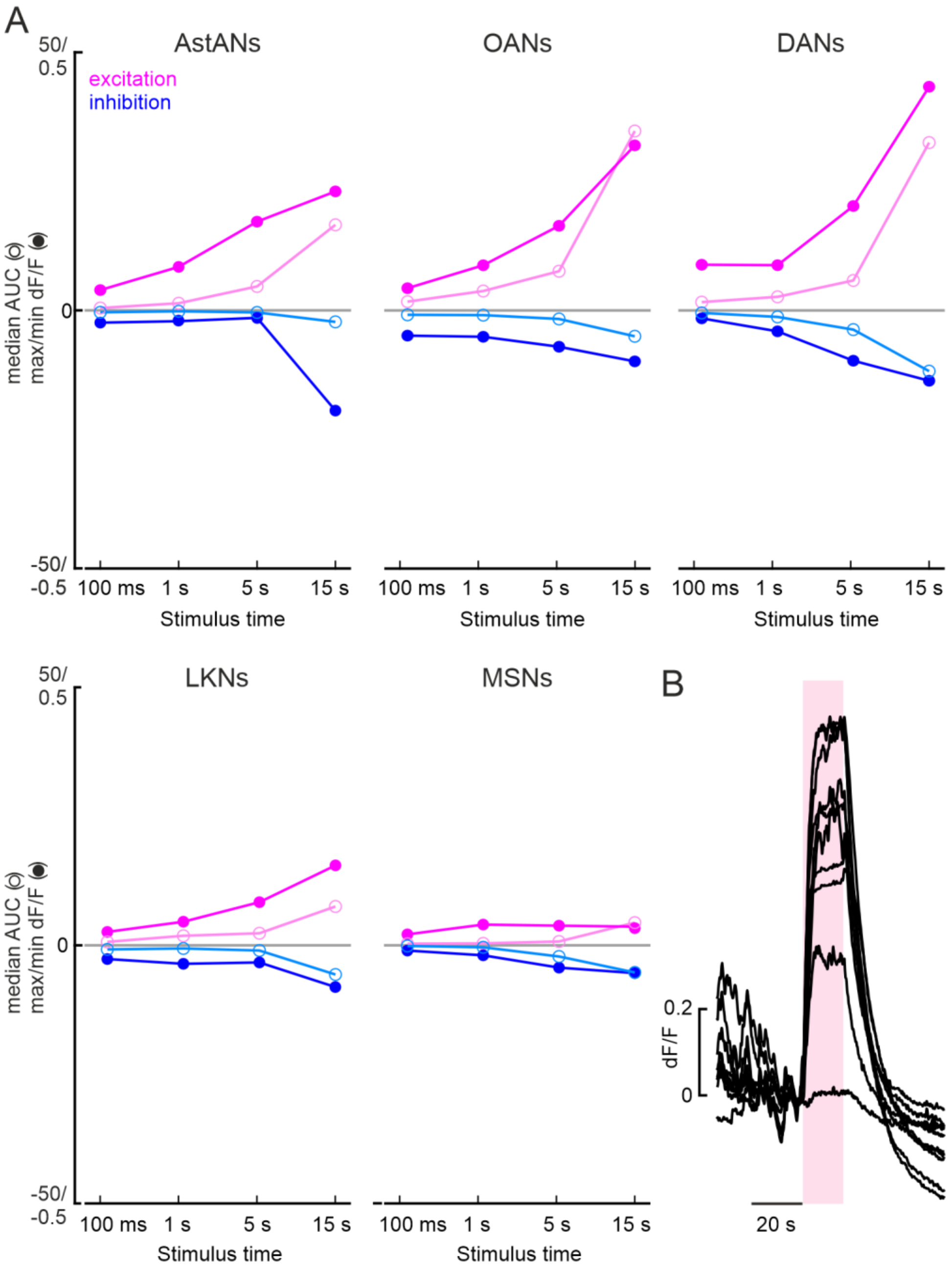


**Figure S9: Comparison of IPC responses to ModN activation for different durations in calcium imaging recordings. A:** Median AUC (empty circles) and maximum and minimum dF/F values (filled circles) for different activation lengths and all ModNs tested. Magenta, excited, blue, inhibited. **B:** Raw traces of 10 IPCs from one fly responding to 15 s AstAN activation revealed strong and robust responses during the whole activation period. After extensive activation, the activity did not return to baseline level, indicating depletion of calcium or deterioration of the IPCs.

**Table S2: G-protein prediction scores for selected receptors in IPCs.** From http://athina.biol.uoa.gr/bioinformatics/PRED-COUPLE2

| Receptor | Isoform | G-protein prediction scores |
|---|---|---|
| AstA-R1 | NP_524700<br>NP_726877 | Gi/o - 0.92<br>Gi/o - 0.92 |
| AstA-R2 | NP_524544<br>NP_001247352<br><br>NP_001263042 | Gi/o - 0.83<br>Gi/o - 0.79<br>Gq/11 - 0.47<br>Gi/o - 0.83 |
| Lkr | NP_647968 | Gq/11 - 0.97 |
| MsR1 | NP_647713<br><br>NP_001261324 | Gi/o - 0.99<br>Gs - 0.32<br>Gi/o - 0.96 |
| MsR2 | NP_647711<br><br>NP_728735<br><br>NP_001261323 | Gi/o - 0.97<br>Gq/11 - 0.48<br>Gi/o - 0.97<br>Gq/11 - 0.48<br>Gi/o - 0.99 |
| TkR86C | NP_524304<br><br>NP_001097741 | Gq/11 - 0.96<br>Gi/o - 0.55<br>Gq/11 - 0.96<br>Gi/o - 0.62 |
| TkR99D | NP_524556<br>NP_001163772<br>NP_001263092 | Gq/11 - 0.93<br>Gq/11 - 0.81<br>Gq/11 - 0.94 |
| sNPF-R | NP_524176<br><br><br>NP_001262086 | Gi/o - 0.99<br>Gq/11 - 0.56<br>G12/13 - 0.38<br>Gi/o - 0.99<br>Gq/11 - 0.56<br>G12/13 - 0.38 |
| Dh31-R | NP_725278<br><br>NP_001260950<br><br><br>NP_001260951 | Gs - 0.83<br>Gi/o - 0.83<br>Gs - 0.88<br>Gi/o - 0.66<br>Gq/11 - 0.32<br>Gi/o - 0.94<br>Gs - 0.91 |
| AkhR | NP_477387<br>NP_723206<br>NP_995639<br>NP_001260149 | Gi/o - 0.99<br>Gi/o - 0.99<br>Gi/o - 0.99<br>Gi/o - 0.99 |
| Dop1R1 | NP_477007<br>NP_001163607<br>NP_001247092<br>NP_001262563<br>NP_001303454 | Gq/11 - 0.62<br>Gq/11 - 0.62<br>Gq/11 - 0.62<br>Gq/11 - 0.57<br>Gq/11 - 0.62 |
| Dop1R2 | NP_733299<br>NP_524548<br>NP_001263072 | Gq/11 - 0.99<br>Gq/11 - 0.99<br>Gq/11 - 0.98 |
| Dop2R | NP_001014759<br>NP_001014757<br><br>NP_001285477<br>NP_001014758<br>NP_001014760<br>NP_001027080 | Gi/o - 0.98<br>Gi/o - 0.54<br>Gq/11 - 0.42<br>Gi/o - 0.83<br>Gi/o - 0.83<br>Gi/o - 0.83<br>Gi/o - 0.72<br>Gq/11 - 0.56 |
| DopEcR | NP_647897<br><br>NP_001014560<br><br>NP_001014559 | Gi/o - 0.97<br>Gq/11 - 0.87<br>Gi/o - 0.97<br>Gq/11 - 0.87<br>Gi/o - 0.97<br>Gq/11 - 0.87 |
| 5-HT1A | NP_725849 | Gi/o - 0.99 |
| 5-HT1B | NP_523789<br>NP_001163201<br>NP_001137708 | Gi/o - 0.64<br>Gi/o - 0.64<br>Gi/o - 0.64 |
| 5-HT2A | NP_524223<br><br>NP_730859<br><br>NP_001163505<br><br>NP_001163506<br><br>NP_001097684 | Gs - 0.92<br>Gq/11 - 0.91<br>Gq/11 - 0.91<br>Gs - 0.89<br>Gs - 0.96<br>Gi/o - 0.93<br>Gq/11 - 0.85<br>Gs - 0.83<br>Gq/11 - 0.85<br>Gs - 0.83 |
| 5-HT2B | NP_001262373<br>NP_649806<br>NP_001287238 | Gs - 0.89<br>Gs - 0.89<br>Gs - 0.89 |
| 5-HT7 | NP_524599<br><br>NP_001263131 | Gs - 0.77<br>Gi/o - 0.42<br>Gs - 0.77<br>Gi/o - 0.42 |
| Oamb | NP_524669<br>NP_732542<br>NP_001262774<br>NP_001303429<br>NP_001262775<br>NP_732541 | Gs - 0.40<br>Gq/11 - 0.86<br>Gq/11 - 0.82<br>Gq/11 - 0.86<br>Gs - 0.40<br>Gq/11 - 0.86 |
| Oct-TyrR | NP_524419<br>NP_001163494 | Gi/o - 0.96<br>Gi/o - 0.96 |
| Octalpha2R | NP_650754<br>NP_001262714<br>NP_001262715 | Gi/o - 0.99<br>Gi/o - 0.99<br>Gi/o - 0.97 |
| Octbeta1R | NP_651057<br>NP_001034064<br><br>NP_001262843<br>NP_001163690 | Gq/11 - 0.56<br>Gq/11 - 0.75<br>Gs - 0.73<br>Gq/11 - 0.56<br>Gs - 0.73<br>Gq/11 - 0.70 |
| Octbeta2R | NP_001034049<br><br>NP_001163596<br><br>NP_001247076<br><br>NP_001247077<br><br>NP_001247078<br><br>NP_001303505 | Gs - 0.60<br>Gi/o - 0.45<br>Gi/o - 0.62<br>Gs - 0.35<br>Gs - 0.60<br>Gi/o - 0.45<br>Gs - 0.60<br>Gi/o - 0.45<br>Gs - 0.60<br>Gi/o - 0.45<br>Gs - 0.60<br>Gi/o - 0.45 |
| Octbeta3R | NP_001034048<br>NP_650210<br><br>NP_001034043<br>NP_001034046 | Gi/o - 0.83<br>Gi/o - 0.88<br>Gs - 0.65<br>Gq/11 - 0.91<br>Gs - 0.98<br>Gi/o - 0.95<br>G12/13 - 0.32 |
